# Neutralization of SARS-CoV-2 spike 69/70 deletion, E484K, and N501Y variants by BNT162b2 vaccine-elicited sera

**DOI:** 10.1101/2021.01.27.427998

**Authors:** Xuping Xie, Yang Liu, Jianying Liu, Xianwen Zhang, Jing Zou, Camila R. Fontes-Garfias, Hongjie Xia, Kena A. Swanson, Mark Cutler, David Cooper, Vineet D. Menachery, Scott Weaver, Philip R. Dormitzer, Pei-Yong Shi

## Abstract

We engineered three SARS-CoV-2 viruses containing key spike mutations from the newly emerged United Kingdom (UK) and South African (SA) variants: N501Y from UK and SA; 69/70-deletion+N501Y+D614G from UK; and E484K+N501Y+D614G from SA. Neutralization geometric mean titers (GMTs) of twenty BTN162b2 vaccine-elicited human sera against the three mutant viruses were 0.81- to 1.46-fold of the GMTs against parental virus, indicating small effects of these mutations on neutralization by sera elicited by two BNT162b2 doses.

## Main

We previously reported that BNT162b2, a nucleoside modified RNA vaccine that encodes the SARS-CoV-2 full length, prefusion stabilized spike glycoprotein (S), elicited dose-dependent SARS-CoV-2–neutralizing geometric mean titers (GMTs) that were similar to or higher than the GMT of a panel of SARS-CoV-2 convalescent human serum samples.^1^ We subsequently reported that, in a randomized, placebo-controlled trial in approximately 44,000 participants 16 years of age or older, a two-dose regimen of BNT162b2 conferred 95% protection against COVID-19.^2^

Since the previously reported studies were conducted, rapidly spreading variants of SARS-CoV-2 have arisen in the United Kingdom (UK), South Africa (SA), and other regions.^3,4^ These variants have multiple mutations in their spike glycoproteins, which are key targets of virus neutralizing antibodies. The emerged spike mutations have raised concerns of vaccine efficacy against these new strains. The goal of this study is to examine the effect of several key spike mutations from the UK and SA strains on BNT162b2 vaccine-elicited neutralization.

Using an infectious cDNA clone of SARS-CoV-2^5^, we engineered three spike mutant viruses on the genetic background of clinical strain USA-WA1/2020 (**Supplementary Fig. 1**). (i) Mutant N501Y virus contains the N501Y mutation that is shared by both the UK and SA variants. This mutation is located in the viral receptor binding domain (RBD) for cell entry, increases binding to the angiotensin converting enzyme 2 (ACE2) receptor, and enables the virus to expand its host range to infect mice.^5,6^ (ii) Mutant Δ69/70+N501Y+D614G virus contains two additional changes present in the UK variants: amino acid 69 and 70 deletion (Δ69/70) and D614G substitution. Amino acids 69 and 70 are located in the N-terminal domain of the spike S1 fragment; deletion of these residues may allosterically change S1 conformation.^6^ The D614G mutation is dominant in circulating strains around the world.^7,8^ (iii) Mutant E484K+N501Y+D614G virus additionally contains the E484K substitution, which is also located in the viral RBD. The E484K substitution alone confers resistance to several monoclonal antibodies.^9,10^ Compared with the wild-type USA-WA1/2020 strain, the three mutant viruses showed similar plaque morphologies on Vero E6 cells (**Supplementary Fig. 2**).

We tested a panel of human sera from twenty participants in the previously reported clinical trial,^1,2^ drawn 2 or 4 weeks after immunization with two 30-μg doses of BNT162b2 spaced three weeks apart (**Supplementary Fig. 3**). All neutralization assays were done with the same 20 sera samples, with the two experiments (as described in **Fig. 1** legend) done at different times. Each serum was tested for neutralization of wild-type USA-WA1/2020 strain and the three mutant viruses by a 50% plaque reduction neutralization assay (PRNT_50_; **Supplementary Tables 1** **and** **2**). All sera showed equivalent neutralization titers between the wild-type and mutant viruses, with differences of ≤ 4 fold (**Fig. 1**). Notably, ten out of the twenty sera had neutralization titers against mutant Δ69/70+N501Y+D614G virus that were twice their titers against the wild-type virus (**Fig. 1b**), whereas six out of the twenty sera had neutralization titers against mutant E484K+N501Y+D614G virus that were half their titers against the wild-type virus (**Fig. 1c**). The ratios of the neutralization GMTs of the sera against the N501Y, Δ69/70+N501Y+D614G, and E484K+N501Y+D614G viruses to their GMTs against the USA-WA1/2020 virus were 1.46, 1.41, and 0.81, respectively (**Supplementary Fig. 4**).

**Figure 1.**
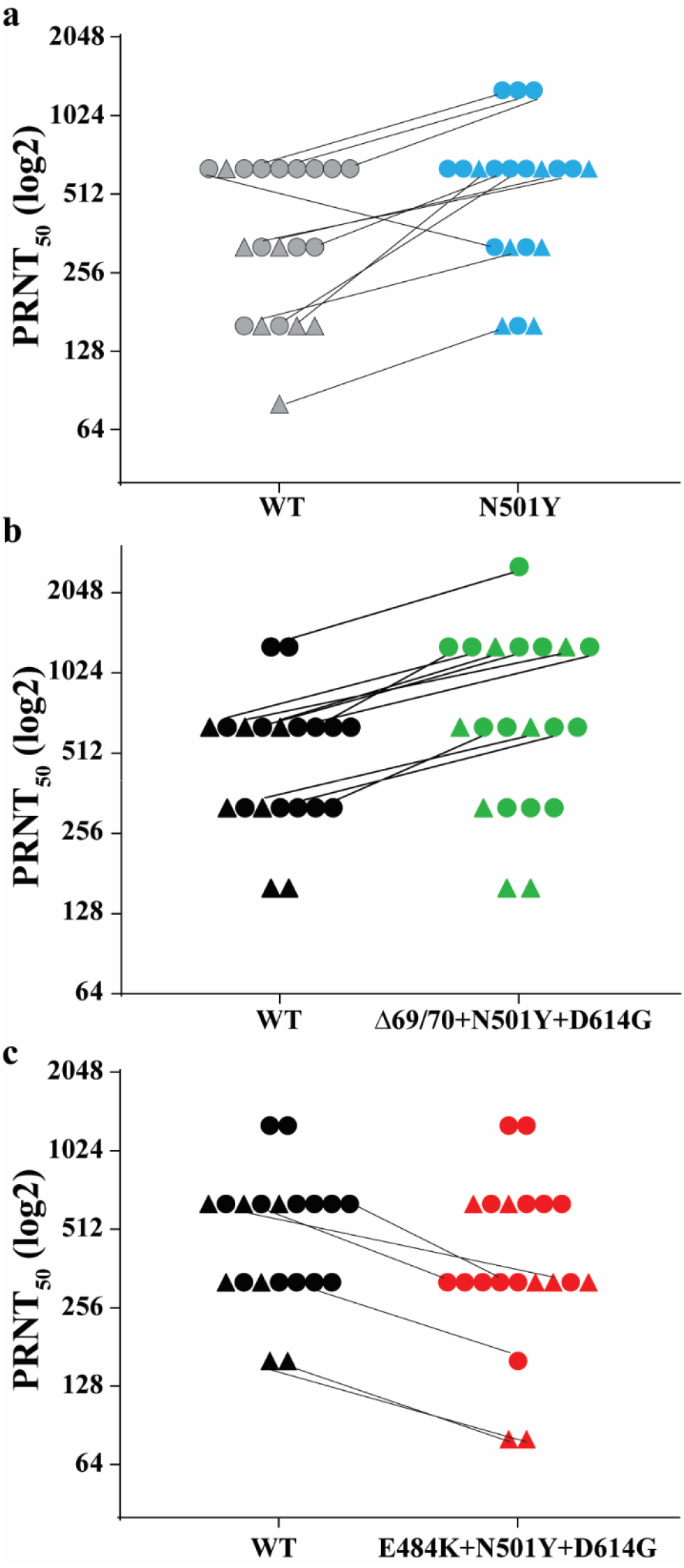
PRNT_50_s of twenty BNT162b2-vaccinated human sera against wild-type (WT) and mutant SARS-CoV-2. **(a)** WT (USA-WA1/2020) and mutant N501Y. **(b)** WT and Δ69/70+N501Y+D614G. **(c)** WT and E484K+N501Y+D614G. Seven (triangles) and thirteen (circles) sera were drawn 2 and 4 weeks after the second dose of vaccination, respectively. Sera with different PRNT_50_s against WT and mutant viruses are connected by lines. Results in (**a**) were from one experiment; results in (**b**) and (**c**) were from another set of experiments. Each data point is the average of duplicate assay results.

Consistent with other recent reports of the neutralization of SARS-CoV-2 variants or corresponding pseudoviruses by convalescent or post-immunization sera,^11,12^ the neutralization GMT of the serum panel against the virus with three mutations from the SA variant (E484K+N501Y+D614G) was slightly lower than the neutralization GMTs against the N501Y virus or the virus with three mutations from the UK variant (Δ69/70+N501Y+D614G). However, the magnitude of the differences in neutralization GMTs against any of the mutant viruses in this study was small (0.81- to 1.41-fold), as compared to the 4-fold differences in hemagglutination-inhibition titers that have been used to signal potential need for a strain change in influenza vaccines.^13^

A limitation of the current study is that the engineered viruses do not include the full set of spike mutations found in the UK or SA variants.^3,4^ Nevertheless, preserved neutralization of N501Y, Δ69/70+N501Y+D614G, and E484K+N501Y+D614G viruses by BNT162b2 vaccine-elicited human sera is consistent with preserved neutralization of a panel of 15 pseudoviruses bearing spikes with other single mutations found in circulating SARS-CoV-2 strains.^14^ The emergence of the common mutation N501Y from different geographical regions, as well as the previously emerged globally dominant D614G mutation, suggest that these mutations may improve viral fitness, as recently demonstrated for the increased viral transmission by the D614G mutation in animal models.^7,15^ The biological functions of N501Y and the other mutations (such as Δ69/70 and E484K) remain to be defined for viral replication, pathogenesis, and/or transmission in animal models. A second limitation of the study is that no serological correlate of protection against COVID-19 has been defined. Therefore, predictions about vaccine efficacy based on neutralization titers require assumptions about the levels of neutralization and roles of humoral and cell-mediated immunity in vaccine-mediated protection. Clinical data are needed for firm conclusions about vaccine effectiveness against variant viruses.

The ongoing evolution of SARS-CoV-2 necessitates continuous monitoring of the significance of changes for vaccine efficacy. This surveillance should be accompanied by preparations for the possibility that future mutations may necessitate changes to vaccine strains. The serological criteria for strain changes of influenza vaccine have been well-accepted.^16^ For COVID-19, such vaccine updates would be facilitated by the flexibility of mRNA-based vaccine technology.

## Methods

### Construction of isogenic viruses

Three recombinant SARS-CoV-2 mutants (N501Y, Δ69/70-N501Y+D614G, E484K+N501Y+D614G in spike protein) were prepared on the genetic background of an infectious cDNA clone derived from clinical strain WA1 (2019-nCoV/USA_WA1/2020)^5^ by following the PCR-based mutagenesis protocol as reported previously^7^. The full-length infectious cDNAs were *in vitro* ligated and used as templates to transcribe full-length viral RNA. Mutant viruses (P0) were recovered on day 2 from Vero E6 cells after electroporation of the *in vitro* RNA transcripts. P1 viruses were harvested as stocks by passaging the P0 virus once on Vero E6 cells. The titers of P1 viruses were determined by plaque assay on Vero E6 cells. The genome sequences of the P1 viruses were validated by Sanger sequencing. The detailed protocol was recently reported^17^.

### Serum specimens and neutralization assay

Serum samples were collected from BNT162b2 vaccinees participating in the phase 1 portion of the ongoing phase 1/2/3 clinical trial (ClinicalTrials.gov identifier: NCT04368728). The protocol and informed consent were approved by institutional review boards for each of the investigational centers participating in the study. The study was conducted in compliance with all International Council for Harmonisation (ICH) Good Clinical Practice (GCP) guidelines and the ethical principles of the Declaration of Helsinki.

The immunization and serum collection regimen are illustrated schematically in **Fig. S3**. A conventional (non-fluorescent) plaque reduction neutralization assay was performed to quantify the serum-mediated virus suppression as previously reported^18^. Briefly, each serum was 2-fold serially diluted in culture medium with the first dilution of 1:40 (dilution range of 1:40 to 1:1280). The diluted sera were incubated with 100 plaque-forming units of wild-type or mutant viruses at 37°C for 1 h, after which the serum-virus mixtures were inoculated onto Vero E6 cell monolayer in 6-well plates. After 1 h of infection at 37°C, 2◻ml of 2% Seaplaque agar (Lonza) in Dulbecco’s modified Eagle medium (DMEM) containing 2% fetal bovine serum (FBS) and 1% penicillin/streptomycin (P/S) was added to the cells. After 2 days of incubation, 2◻ml of 2% Seaplaque agar (Lonza) in DMEM containing 2% FBS, 1% P/S and 0.01% neutral red (Sigma) were added on top of the first layer. After another 16 h of incubation at 37°C, plaque numbers were counted. The minimal serum dilution that inhibits 50% of plaque counts is defined as the 50% plaque reduction neutralization titer (PRNT_50_). Each serum was tested in duplicates. The PRNT_50_ assay was performed at the biosafety level-3 facility with the approval from the Institutional Biosafety Committee at the University of Texas Medical Branch.

### Statistics

No statistics were performed in the study.

### Data availability

The data that support the findings of this study are available from the corresponding authors upon request.

## Acknowledgments

Supported by Pfizer and BioNTech. We thank colleagues at Pfizer, BioNTech, and UTMB for helpful discussions and support during the study. We thank the Pfizer-BioNTech clinical trial C4591001 participants, from whom the post-immunization human sera were obtained. We thank the many colleagues at Pfizer and BioNTech who developed and produced the BNT162b2 vaccine candidate. P.-Y.S. was supported by NIH grants AI142759, AI134907, AI145617, and UL1TR001439, and awards from the Sealy & Smith Foundation, Kleberg Foundation, the John S. Dunn Foundation, the Amon G. Carter Foundation, the Gilson Longenbaugh Foundation, and the Summerfield Robert Foundation.

## Author contributions

Conceptualization, X.X., V.D.M., S.W., P.-Y.S.; Methodology, X.X., Y.L., J.L., J.Z., C.R.F.G., H.X., P.-Y.S; Investigation, X.X., Y.L., J.L., J.Z., C.R.F.G., H.X., K.A.S., D.C., P.R.D., P.-Y.S; Resources, M.C., D.C., P.R.D., P.-Y.S; Data Curation, X.X., Y.L., J.L., J.Z., C.R.F.G., P.-Y.S; Writing-Original Draft, X.X., P.-Y.S; Writing-Review & Editing, X.X., P.R.D., P.-Y.S.; Supervision, X.X., M.C., D.C., P.R.D., P.-Y.S.; Funding Acquisition P.-Y.S.

## Supplementary information

**Supplementary Figure 1.**
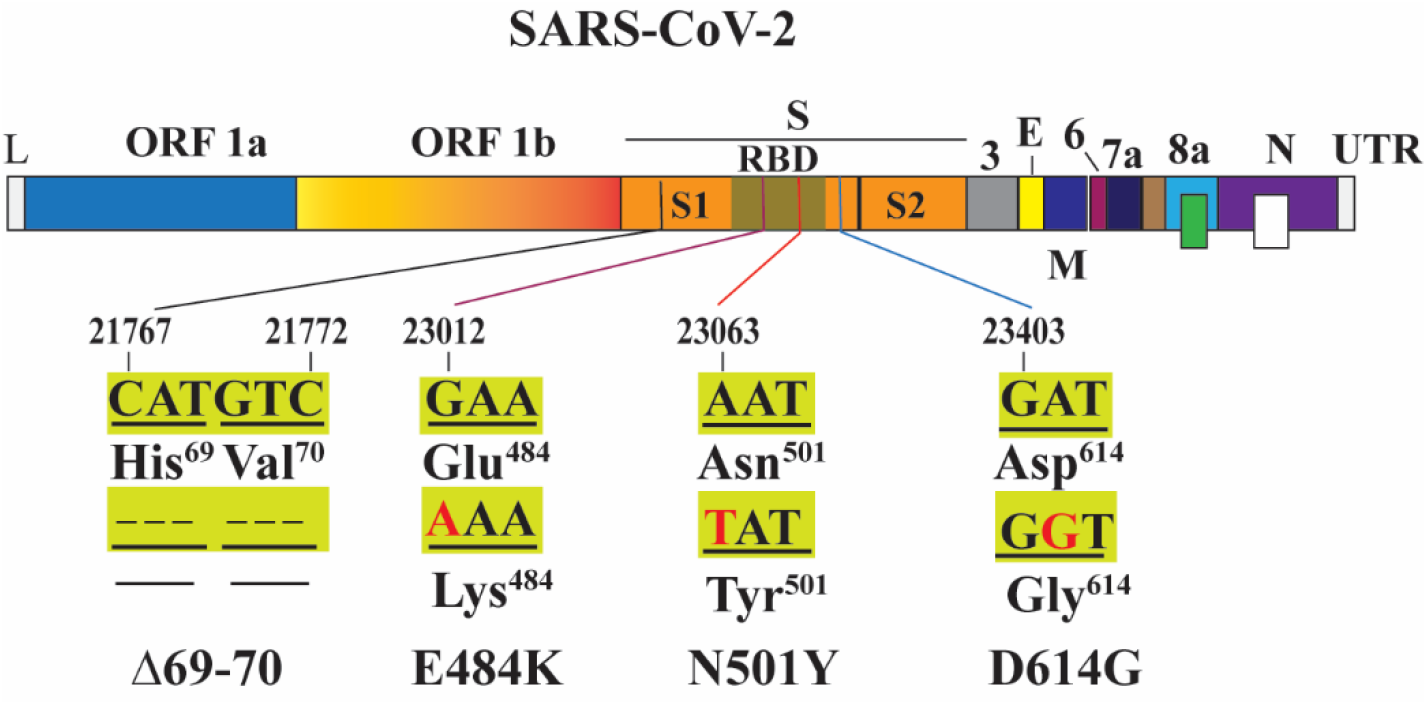
Engineered mutations. Nucleotide and amino acid positions are indicated. Deletions are depicted by dotted lines. Mutant nucleotides are in red. L, leader sequence; ORF, open reading frame; RBD, receptor binding domain; S, spike glycoprotein; S1, N-terminal furin cleavage fragment of S; S2, C-terminal furin cleavage fragment of S; E, envelope protein; M, membrane protein; N, nucleoprotein; UTR, untranslated region.

**Supplementary Figure 2.**
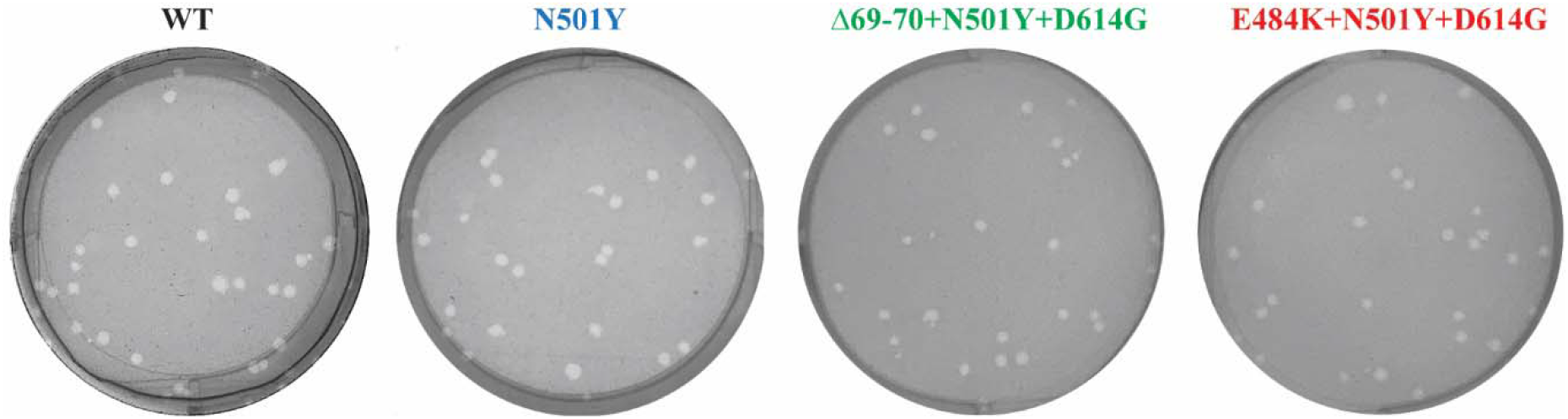
Plaque morphologies of WT (USA-WA1/2020), mutant N501Y, Δ69/70+N501Y+D614G, and E484K+N501Y+D614G SARS-CoV-2s on Vero E6 cells.

**Supplementary Figure 3.**
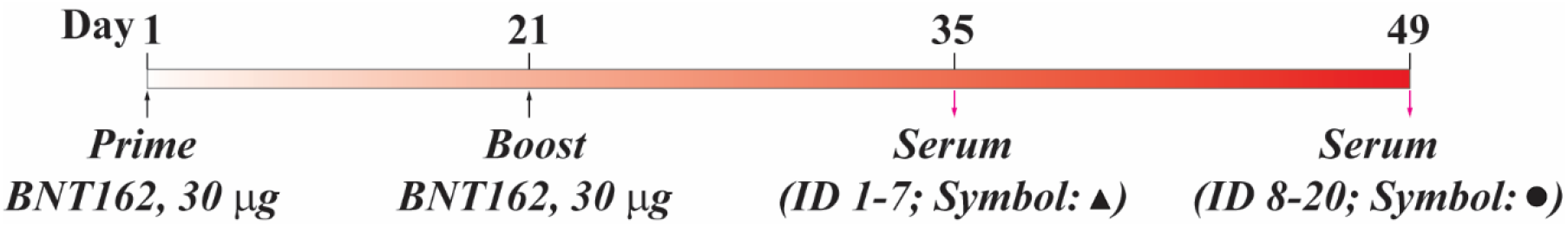
Scheme of the BNT162 vaccination and serum sampling.

**Supplementary Figure 4.**
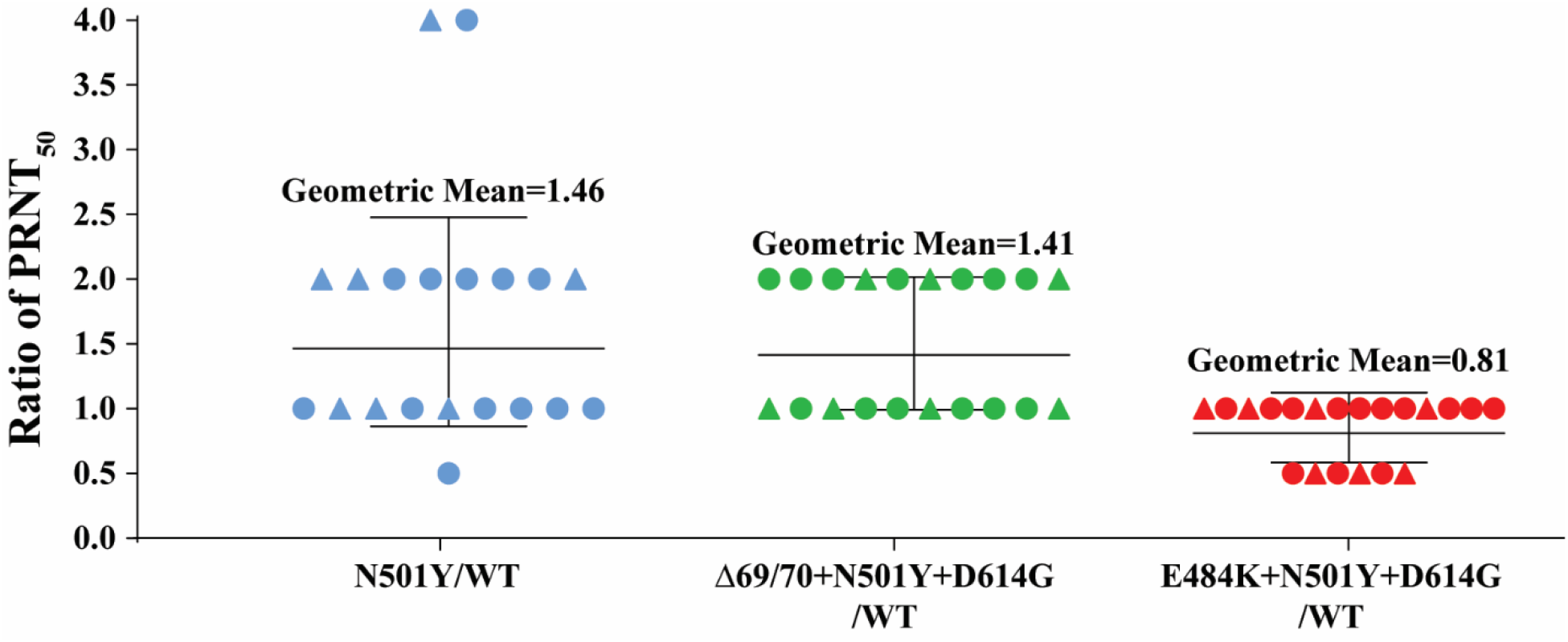
Ratios of neutralization GMTs against mutant viruses to GMTs against WT virus. Triangles represent sera drawn two weeks after the second dose of vaccination; circles represent sera drawn four weeks after the second dose of vaccination.

**Supplementary Table 1.** PRNT_50_s of twenty BNT162b2 post-immunization sera against wild-type (USA-WA1/2020) and mutant N501Y SARS-CoV-2s.

| Serum ID | PRNT<sub>50</sub> |  | PRNT<sub>50</sub> ratio<br>(N501Y/WT) |
| --- | --- | --- | --- |
|  | WT | N501Y |  |
| 1 | 160 | 640 | 4 |
| 2 | 160 | 320 | 2 |
| 3 | 320 | 640 | 2 |
| 4 | 80 | 160 | 2 |
| 5 | 160 | 160 | 1 |
| 6 | 320 | 320 | 1 |
| 7 | 640 | 640 | 1 |
| 8 | 160 | 160 | 1 |
| 9 | 640 | 640 | 1 |
| 10 | 640 | 1280 | 2 |
| 11 | 160 | 640 | 4 |
| 12 | 320 | 320 | 1 |
| 13 | 640 | 1280 | 2 |
| 14 | 640 | 320 | 0.5 |
| 15 | 320 | 640 | 2 |
| 16 | 320 | 640 | 2 |
| 17 | 640 | 640 | 1 |
| 18 | 640 | 1280 | 2 |
| 19 | 640 | 640 | 1 |
| 20 | 640 | 640 | 1 |

**Supplementary Table 2.**
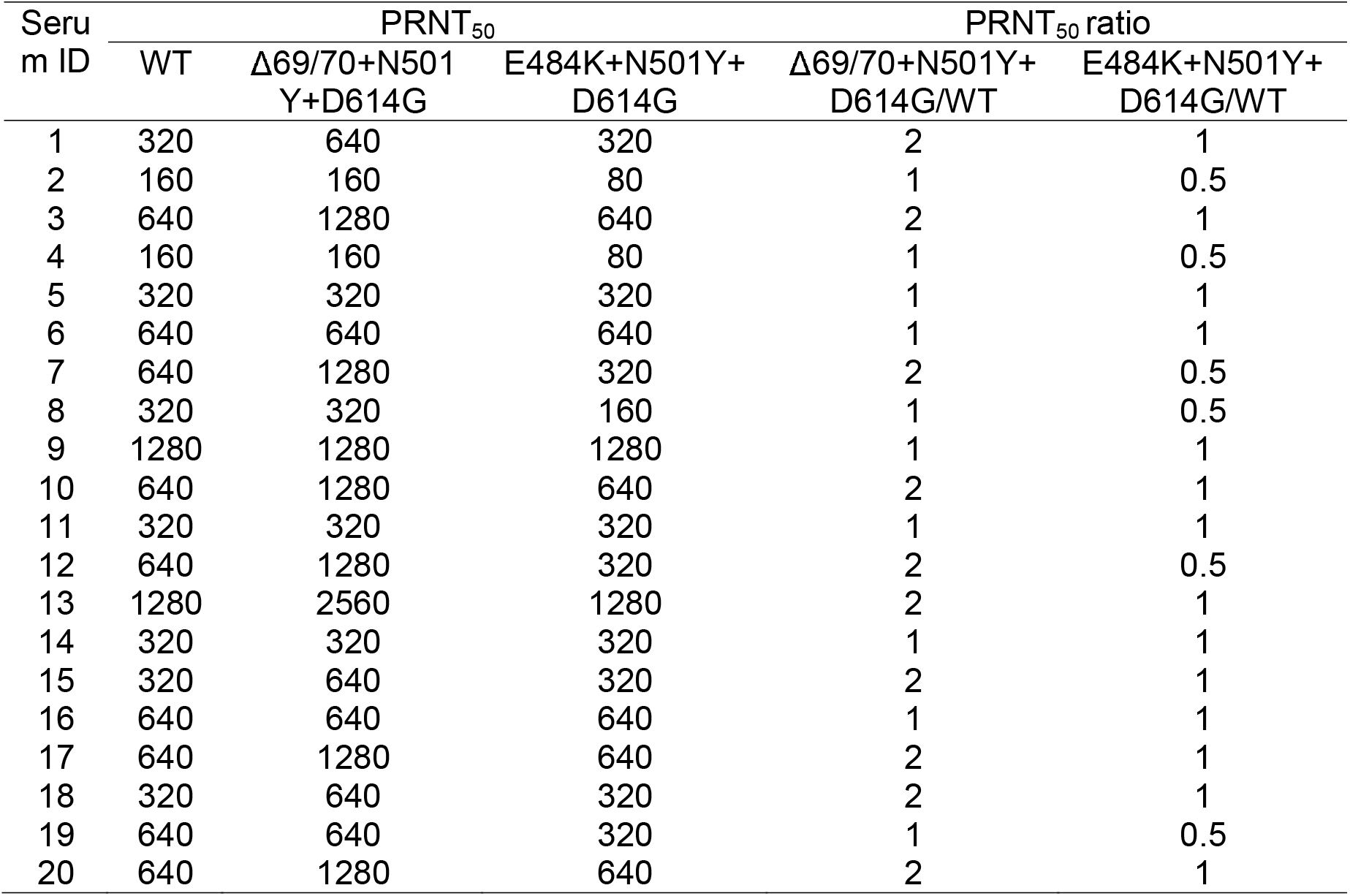
PRNT_50_s of twenty BNT162b2 post-immunization sera against wild-type (USA-WA1/2020), Δ69/70+N501Y+D614G, and E484K+N501Y+D614G SARS-CoV-2s.

